# Computational design of DNA binding domain-retained fusion proteins and virtual screening against FDA-approved drugs

**DOI:** 10.1101/2023.05.05.539610

**Authors:** Himansu Kumar, Pora Kim

## Abstract

Even though the transcription factors (TFs) are not regarded as good drug targets, mutated or dysregulated TFs can be a unique class of drug targets. Specifically, the TF fusion protein, which is the translated structural variants including TFs may affect downstream to promote tumorigenesis. To date, we lack the fusion protein sequence information and 3D structure information in identifying the potential drugs of fusion proteins. In this study, we predicted the 3D structures of 732 transcription factor fusion proteins (TFFPs). For the top five most frequent TFFPs, we performed the virtual screening across the FDA-approved drugs. Our study will provide an initial platform to develop novel therapeutic targets in the transcription factor fusion proteins.

## INTRODUCTION

Fusion genes have been recognized as important biomarkers and therapeutic targets in a variety of cancers. With the huge accumulated structural variant data sets, to help better understand the function of the coding structural variants, previously we developed a unique resource, named FusionGDB (Fusion Gene annotation DataBase v1 and v2) providing intensive functional annotation of ∼ 126K human fusion genes using bioinformatics and deep learning approaches including ∼ 83K full-length fusion transcript sequences and ∼ 43K fusion amino acid sequences based on the multiple breakpoints and gene isoforms. Recently, we analyzed the functional annotation of ∼ 43K fusion proteins, predicted the whole 3D structures of 2.3K and 1.3K fusion proteins from 2.3K recurrent fusion genes and 266 manually curated fusion genes, and performed the virtual screening of 1.3K fusion proteins of 266 manually curated fusion genes across the FDA-approved drugs. This new knowledge on human cancer fusion proteins can be accessed through the website of FusionPDB.

Multiple mechanisms of action can be used to categorize fusion genes, including the signal transduction with control of kinase fusion genes, target gene activation of the transcription factor fusion genes, lost protein-protein interaction, avoidance of miRNA regulation, up-regulated downstream effectors, and gain/loss of cellular regulatory subunit ^1^. The kinase fusion genes got the most attention so far since the success of the development of imatinib targets the BCR-ABL1 kinase fusion protein. Out of 500 human kinomes, 485 kinases are targeted by the drugs from IUPHAR target information. On the contrary, only 80 out of 1429 human TFs were targeted by drugs so far from IUPHAR target information. However, since the aberrant DNA binding activity could cause downstream effects on tumorigenesis, the transcription factor fusion proteins may have a chance to affect their target gene regulation. For example, the TMPRSS2-ERG fusion gene has over 50% of frequency in prostate cancer patients. TMPRSS2, which is a prostate tissue-specific and androgen-response gene, enhanced the function of proto-oncogene (ERG), which has an ETS DNA binding domain, so results in the overexpression of target genes. Androgen ablation therapies are used to treat these prostate patients ^2^. Similarly, if a transactivation domain of EWSR1 is fused with a DNA-binding domain of FLI1, this fusion protein can recruit transcriptional coregulatory elements, resulting in target gene activation ^1^. TK216 is under the clinical trial for Ewing sarcoma patients targeting the DNA-binding domain of EWSR1-FLI1 fusion ^3^.

In this study, for 731 TF fusion proteins of 127 manually curated TF fusion genes, we checked the protein functional features’ retention, predicted the best active sites based on the whole 3D structures, and performed virtual screening to check the approved drugs that may have potential interaction. We chose the top five most frequent TFFPs; TMPRSS2-ERG, KMT2A-AFF1, RUNX1-RUNX1T1, PML-RARA, and EWSR1-FLI1. For these fusion proteins, we predicted potentially interacting FDA-approved drugs. In the investigation of the top 10 interacting drugs of these fusion proteins in their cancer-type context, we suggest potentially interacting drugs. As the first systematic study providing the whole 3D protein structures, a landscape of human transcription factor fusion proteins, and potentially interacting small molecules, our study will provide an initial platform for the drug development of the fusion proteins in cancers.

## RESULTS

### 732 curated human transcription factor fusion proteins and domain retention search

Out of 1,597 manually curated human fusion genes from the ChimerKB4 of ChimerDB4 ^4^, 1,007 fusion genes have the designated fusion breakpoint information. After checking the open reading frames, we had 272 in-frame fusion genes. After transcription of these fusion genes considering multiple breakpoints and multiple gene isoforms, we made 2,485 full-length fusion transcript sequences. Inputting these fusion transcript sequences into the ORFfinder and choosing the longest ORFs, we finally had 1,267 translated fusion amino acid sequences. Interestingly, out of 266 manually curated in-frame fusion genes, 47.7% (127/266*100) included the transcription factors as one of the partners in the fusion genes. This percentage was higher in the number of fusion amino acid sequences at 57.77% (732/1267*100). (Figure 1A) The fusion gene information and their sequences (full-length fusion transcript sequences and fusion amino acid sequences) are in Supplementary Table S1. For all 732 TF fusion proteins, we predicted the full 3D structures using AlphaFold2 ^5^ and virtually screened the potentially interacting small molecules based on the FDA-approved drug library. As the benchmark data, we chose the top five frequently expressed TF fusion genes (Figure 1A). After checking the retention of the DNA binding domain of individual TFs of TFFGs, we constructed the TF fusion gene network using the DNA binding domain retained cases only. As shown in Figure 1B, the KMT2A, a transcription factor or a coactivator, saved as a TF in TRANSFAC, was involved in the fusion gene with many other partner genes. Even though EWSR1 was not a transcription factor, it was involved in many fusion genes with multiple TFs. ETV6 transcription factor was involved in multiple fusion genes with kinases. To see the impact of individual TFs in pan-cancer fusion genes, we calculated the fusion gene assessment scores of 127 TFs that were involved in the formation of 127 TFFGs (732 TFFPs) (Figure 1C). The Major Active Iso-fusion Index (MAIIs) shows the number of samples reported of all possible combinations of fusion genes from multiple cancers, multiple partner genes, and multiple breakpoints of each TF. The positive values describe that a TF has a high sample frequency compared to the number of all possible combinations of formation of fusion genes. KMT2A, ERG, and ETV6 seem actively involved in the TFFGs in pan-cancer.

**Figure 1.**
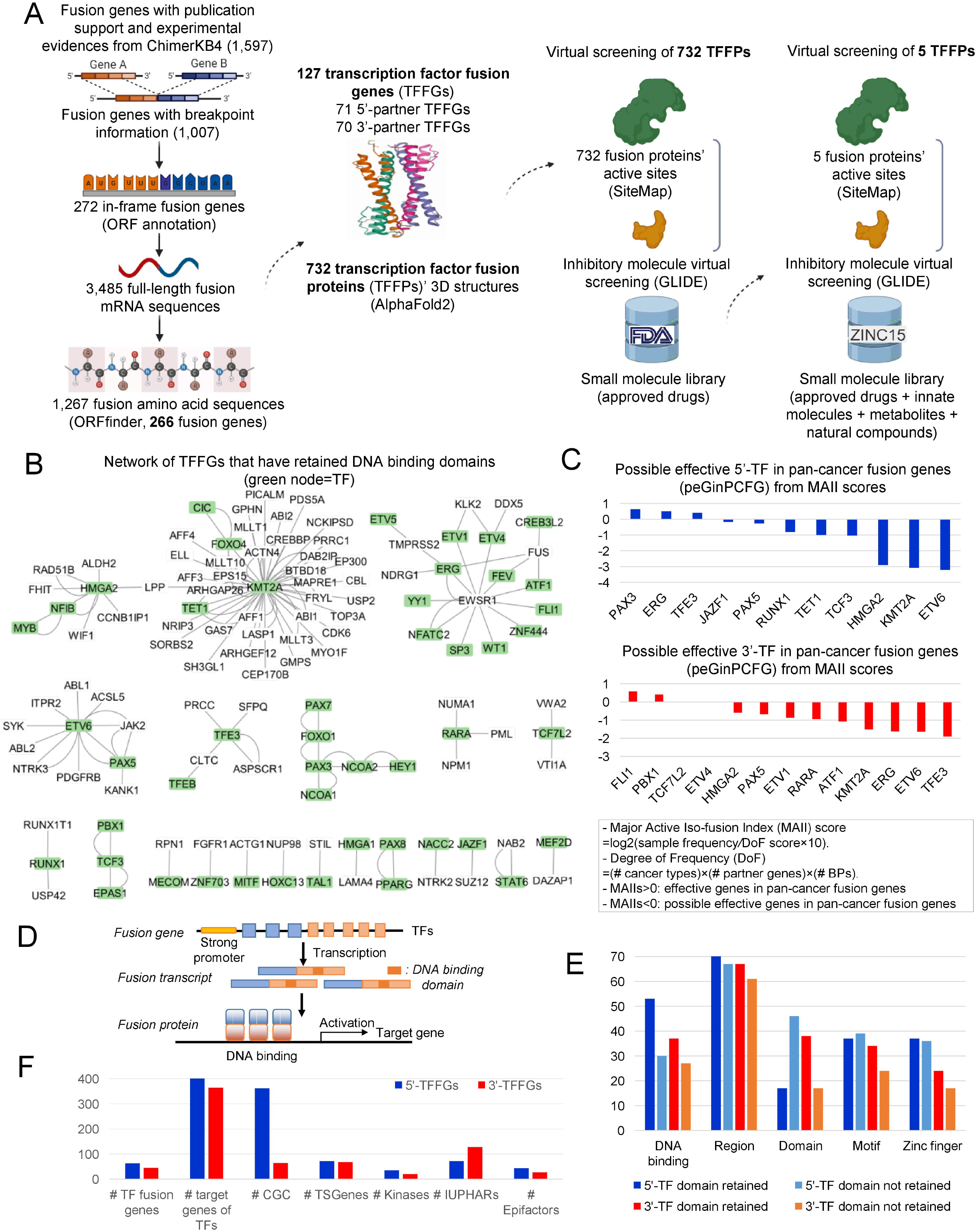
Overview of our analysis pipeline and a network of TFFGs with assessed partner genes in pan-cancer fusion genes. (A) Flowchart of our study. We first annotate the open reading frames of ∼ 1007K human fusion genes from ChmerKB4. For 266 in-frame fusion genes, we made ∼ 3,485 full-length fusion transcript sequences considering multiple breakpoints and gene isoforms. Using ORFfinder, we made 1,267 fusion amino acid sequences. Out of 226 manually curated in-frame fusion genes, there were 127 transcription factor fusion genes, which can produce 732 transcription factor fusion proteins. For these TFFPs and the top five most frequent TFFPs, we performed virtual screening versus FDA-approved drugs and 320K small molecule libraries, respectively. (B) For 127 TFFGs, we investigated the DNA binding domain retention. Using the DNA binding domain-retained fusion genes, we constructed a gene-gene network. The green nodes denote the TFs. (C) Using our calculation of the assessment of individual partner genes in pan-cancer fusion genes, two bar plots show the sorted possible effective 5’- and 3’-partner TFs. (D) Conceptual image of the general tumorigenic mechanism of TFFGs through the target gene activation while the TFFGs retain the DNA binding domain. (E) Distribution of important gene groups and UniProt protein functional major features in the 5’-retained/not-retained and 3’-retained/not-retained fusion genes. (F) Statistics of the target genes of 5’- and 3’-TF fusion genes.

Figure 1D shows the conceptual image of the general mechanism of action of the transcription factor fusion genes. If a fusion protein retains the DNA binding domain (DBD) binding to its target genes’ promoter regions. If the target genes have roles in tumorigenesis, then this TF fusion protein can contribute to tumorigenesis. We investigated the major functional features that may include the DNA binding domain features out of all 39 UniProt protein functional features of 735 TF fusion proteins ^6^ (Figure 1E). Then, ‘DNA binding’ and ‘Zinc finger’ protein functional feature was more retained in the 5’-partner TFs compared to the 3’-TFs. On the other hand, in the 3’-partner TFs, ‘Domain’ protein functional feature was more retained compared to 5’-TF. 63 and 45 TF fusion genes that have TFs in their 5’- and 3’-partner genes, there were 450 and 364 known target genes, respectively, according to the TF-target pair information of the TRRUST database (v2) ^7^. Interestingly, out of 450 target genes of the 5’-TFs, there were 362 cancer gene census (CGC) genes, and out of 364 target genes of the 3’-TFs, there were 128 IUPHAR drug targets (Figure 1F and Supplementary Table S2). This may reflect the importance of the drug development of the TF fusion proteins in human cancer.

### Prediction of the whole 3D structures of the transcription factor fusion proteins

We predicted the 3D structures of 732 transcription factor fusion proteins using AlphaFold2 ^5^. The outputs of AlphaFold2 provide the 3D structure PDB file with the reliability scores of a per-residue confidence score (pLDDT scores). Figure 2A shows the distribution of the number of fusion proteins across the pLDDT averaged score range. Overall, the 3D structures of 732 TFFPs have good pLDDT scores, which are larger than 50. Figure 2B shows the distribution of pLDDT scores across protein sequence length of the 5’-partner wild-type proteins, fusion proteins, and 3’-partner wild-type proteins. In these five top most frequent fusions, individual DNA binding domains were retained, as highlighted in the yellow rectangle in the protein structure. Figure 2C shows the superimposed structures of predicted fusion proteins by AlphaFold2 and the known structures of individual transcription factor proteins from PDB ^8^. The calculated RMSD values were less than 2.0 Å, corresponding to reasonable docking solutions. Figure 2D shows the theoretically possible secondary structure information by the Ramachandran plots of these five fusion proteins. It shows that no more than 2% of residues reside in the disallowed region.

**Figure 2.**
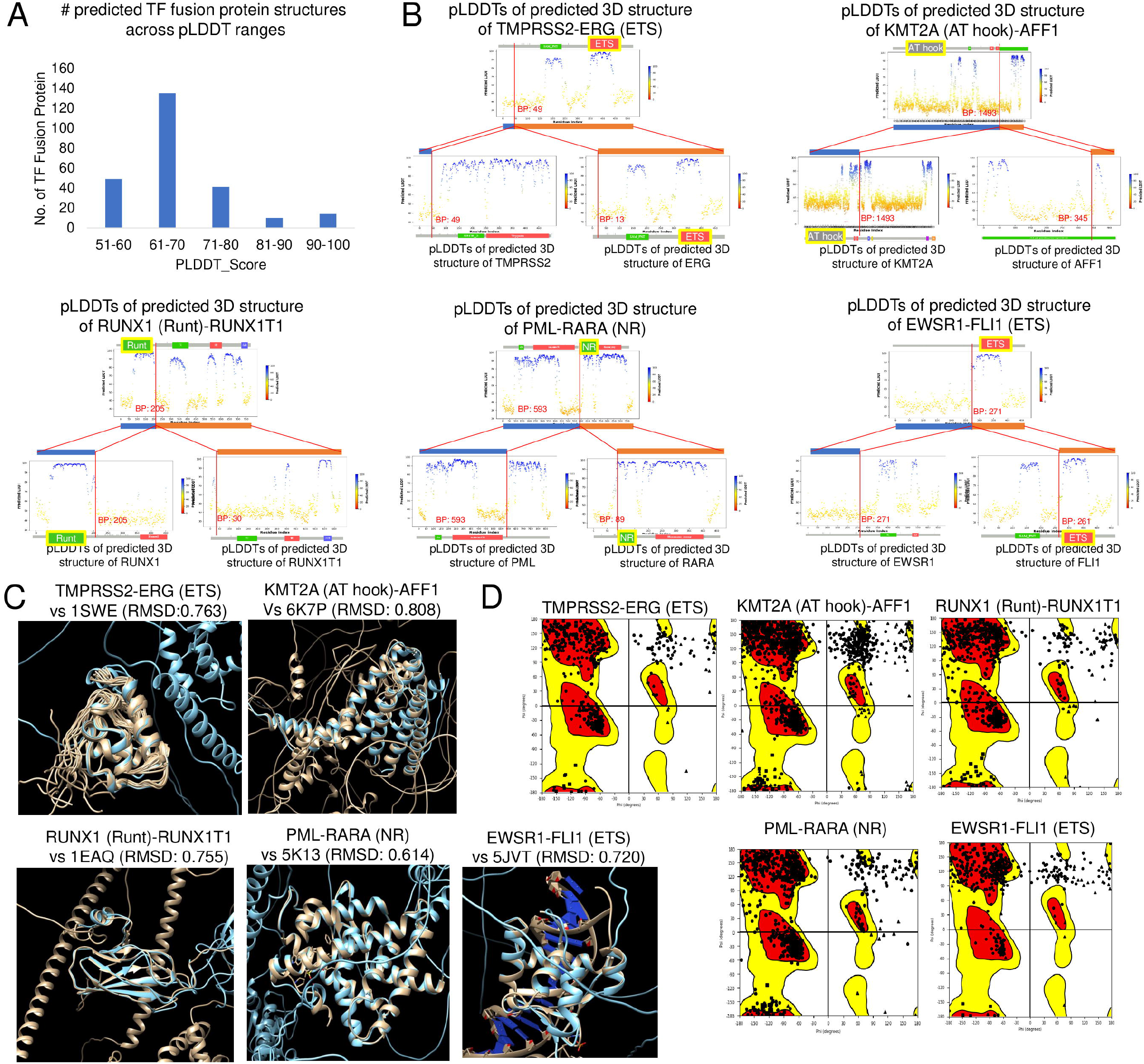
Prediction of 3D structures of the top five most frequent TFFPs. We predicted the 3D structures of fusion proteins using AlphaFold2. (A) The distribution of pLDDT scores across the fusion protein sequence length and the one for 5’-partner and 3’-partner wild-type proteins. (B) pLDDT score distribution of all 732 predicted whole 3D structures of TF fusion proteins. (C) Superimposed structures of five predicted TFFPs by AlphaFold2 and known structure of individual TFs from PDB. The blue color is the predicted structure by AlphaFold2, and grey indicates the reference PDB. These fusion proteins were predicted based on the most frequent. RMSD between the predicted structure and PDB reference were calculated. (D) Ramachandran plots of the predicted whole 3D structures of five TFFPs.

### Virtual screening between 732 TFFPs and FDA-approved drugs

First, we chose the best active sites from the ranked scores of SiteMap based on the whole 3D structures of TFFPs, then checked the location of the best active sites whether in the major functional domain regions. Based on these best active sites, we performed the virtual screening between 732 TFFPs and the FDA-approved drugs, which was aimed to avoid the off-target effects and to identify potential drug repurposing. Among all virtual screening lists, for the top ten FDA-approved drugs interacting with TFFPs, we counted the number of fusion genes interacting with these molecules per drug type (Figure 3A). Then, the most frequent types of drugs were the inhibitors, antagonists, and agonists. To provide a more intuitive landscape of the interactions between individual transcription factors and interacting drugs, we built a TF-drug network (Figure 3B). To make this network, we chose the TFs that are involved in the formation of fusion proteins with more than ten different partners and the top ten interacting FDA-approved drugs with these fusions. From the top left, sorted transcription factors, including NFIB, ERG, MYB, TCF7L2, and KMT2A, have high interaction with more than ten FDA-approved drugs targeting more than ten different TFs from the screening of 732 TFFPs. The top five most frequently listed from the screening were Temozolomide, Chlorthalidone, Mitoxantrone, Ceftolozane, and Clavulanic acids. The followings are the summary of these molecules from DrugBank^9^. Temozolomide is an alkylating agent used to treat glioblastoma multiforme and refractory anaplastic astrocytoma. Chlorthalidone is a diuretic used to treat hypertension or edema caused by heart failure, renal failure, hepatic cirrhosis, estrogen therapy, and other conditions. Mitoxantrone is a chemotherapeutic agent used for the treatment of secondary progressive, progressive relapsing, or worsening relapsing-remitting multiple sclerosis. Ceftolozane is a cephalosporin antibiotic used to treat complicated intra-abdominal infections in combination with metronidazole, complicated urinary tract infections, and hospital-acquired pneumonia. Clavulanic acid is a beta-lactamase inhibitor used to enhance the effectiveness of beta-lactam antibiotics.

**Figure 3.**
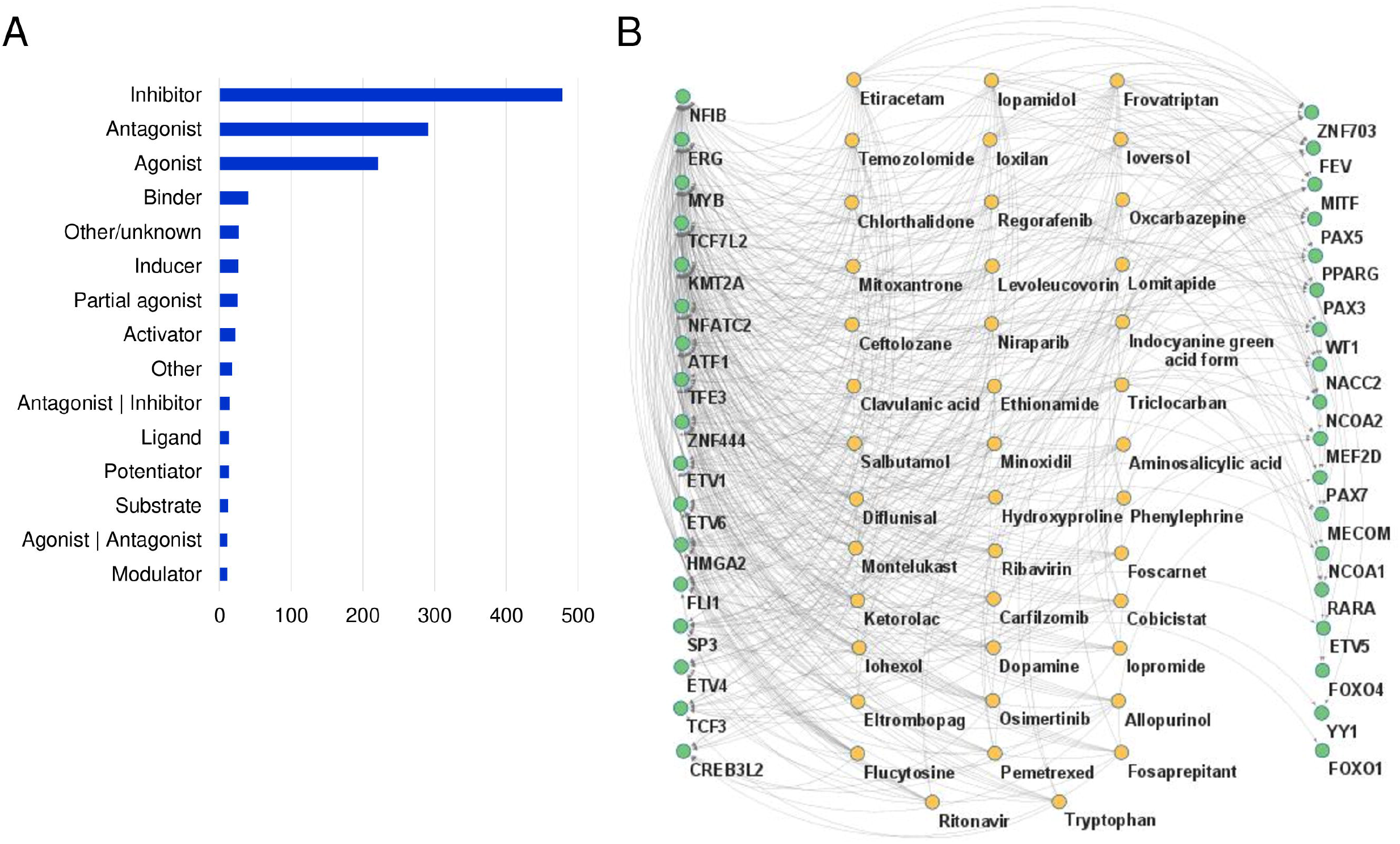
Top 10 interacting FDA-approved drugs with ∼ 750 TFFPs. (A) The number of fusion genes interacting with the top 10 FDA-approved drugs of ∼ 750 TFFPs per drug type. (B) Top 10 FDA-approved drugs interacting with ∼ 750 TFFPs targeting more than 10 different partner genes in TF fusion genes. (C) Top 10 individual cancer-associated drugs through virtual screening using different active sites of the top five most frequent TFFPs.

### Comparison of the virtual screening results based on the whole fusion protein 3D structure and TF domain structure

Active sites are very critical to screen the interacting molecules. To date, the active sites were mainly predicted on the functional domain only due to the lack of available whole 3D structures of fusion proteins. This strategy can be failed to identify the fusion protein specifically interacting small molecules and hard to have knowledge of the best active sites in the whole 3D structure context. It also can lead to listing the weak interacting small molecules than strong binding due to not considering the whole 3D structure context. In this study, we predicted the active sites of 732 TFFPs based on their whole 3D structures using the SiteMap of the Schrödinger package ^10^. To see how different the virtual screening results are using the whole structure-based active sites versus the known biological functional domain, we made two types of active sites and performed the virtual screening with FDA-approved drugs. For further analysis of the identification of the potential interaction molecules, we chose the top five most frequent TFFGs from their sample frequency. Those TFFGs are TMPRSS2-ERG, KMT2A-AFF1, RUNX1-RUNX1T1, PML-RARA, and EWSR1-FLI1, which include five transcription factors (ERG, KM2TA, RUNX1, RARA, and FLI1) as one of the fusion partner genes in the prostate cancer, acute lymphoblastic leukemia (ALL), acute myeloid leukemia (AML), acute promyelocytic leukemia (APL), and Ewing sarcoma, respectively. Figure 4A shows detailed information on the known DNA binding domains and active domains of the top five most frequent TFFPs. Figure 4B shows the best active sites based on the whole 3D structures (Figure 4B) sorted by the SiteMap scores (SiteScore). In TMPRSS2-ERG, DNA binding domain and active domain lie between 337 to 347 and 299 to 317 amino acids, respectively, from UniProt. The SiteMap predicted active sites were from 269 to 318 amino acid residues. Figure 4C shows the distribution of these different active sites in 3D for comparison. For the known biological domains, we made longer amino acids covering both transcription factor domains (DNA binding domain and active domain). For example, for TMPRSS2-ERG, we set the active sites as 290 to 400. We made the grid around these residues for molecular docking with the help of the Receptor Grid Generation Panel of Schrodinger. For other four fusion proteins, we similarly set the active sites and performed the virtual screening and docking.

**Figure 4.**
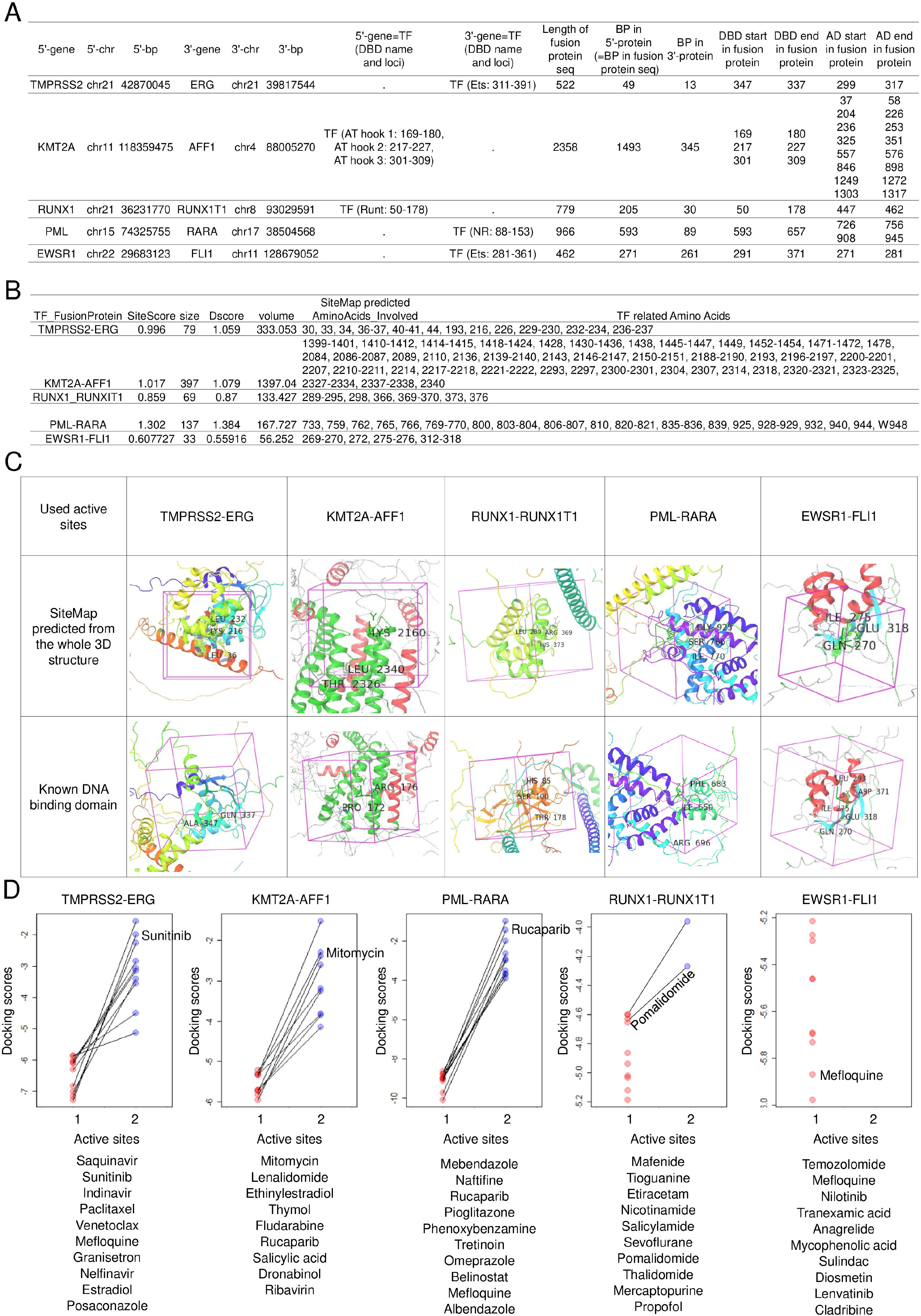
Active site details of the top five most frequent TFFPs. (A) Known DNA binding domain (DBD) and active domain (AD) information across fusion protein sequence. (B) Predicted active site information from SiteMap across fusion protein sequence. (C) Active sites in 3D.

Supplementary Tables S3A and S3B show the top 10 FDA-approved drugs interacting between the whole 3D structure-based active sites and the known biological domains of five TFFPs, respectively. The result was very different between the virtual screening of different active sites. There was only one drug commonly listed in the top 10 drugs from these two different screenings in the same fusion protein. Based on the cancer context of five TFFPs, we chose the potential repurposing drugs in Table 1. As shown in Figure 4D, overall the chosen cancer-associated drugs had stronger interaction in the whole 3D structure-based predicted active sites than in the known TF domains. Specifically, most of the drugs of the RUNX1-RUNX1T1 and EWSR1-FLI1 did not form any interaction from the screening based on the known TF domains. Supplementary Table S3C shows the docking scores between the virtual screening results based on our predicted best active sites and known domains for the chosen best-candidate drugs.

**Table 1.**
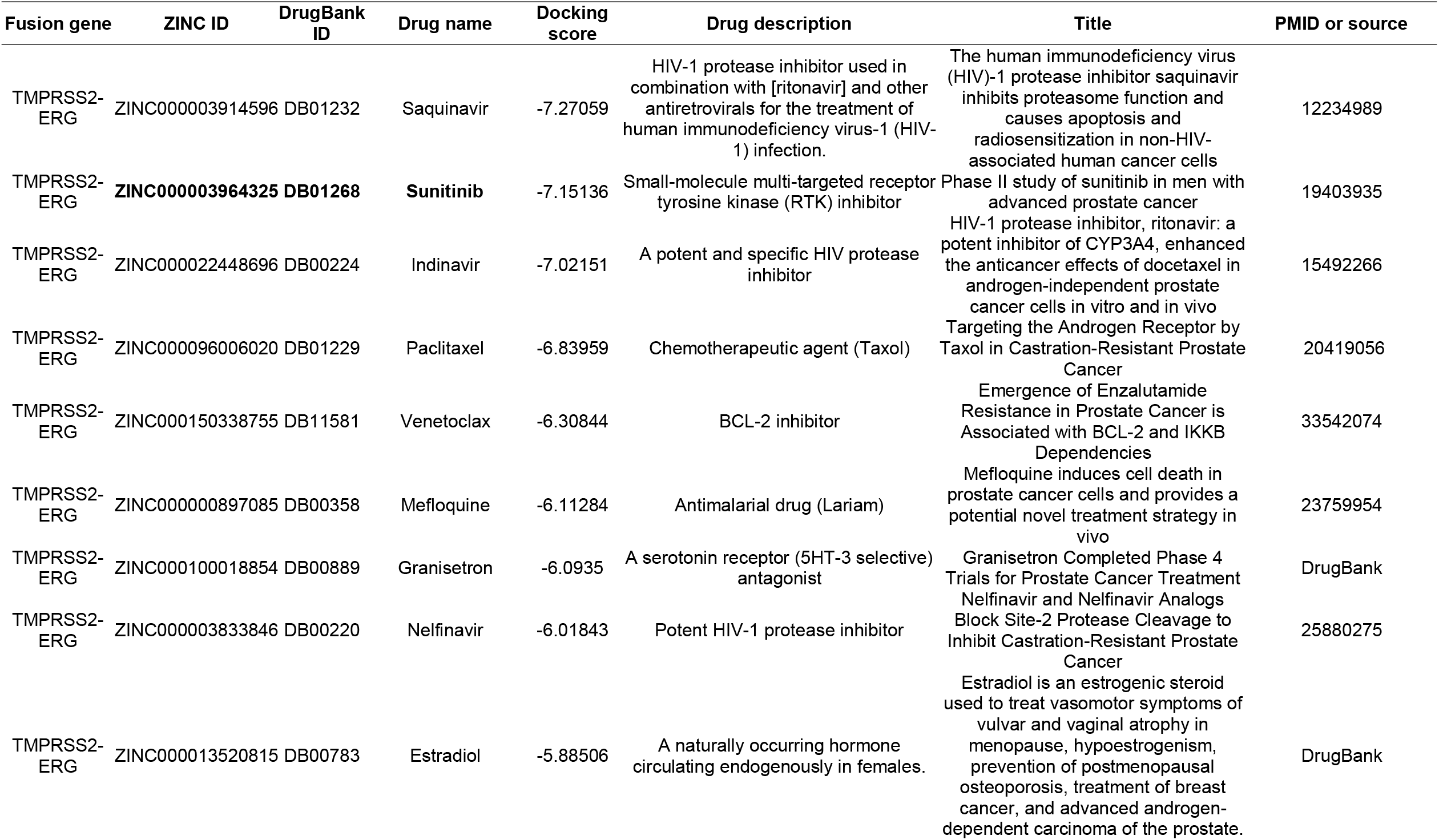
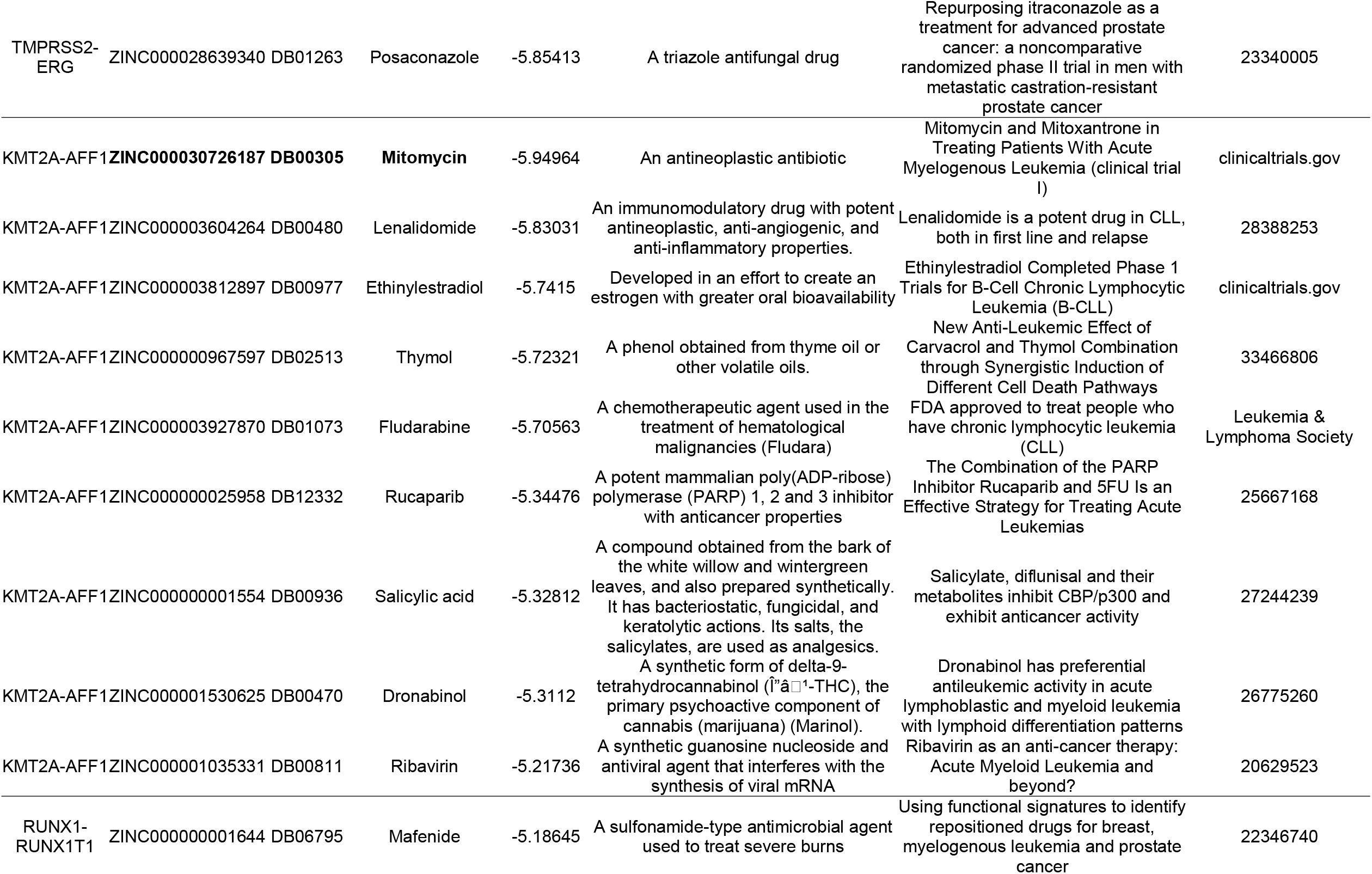
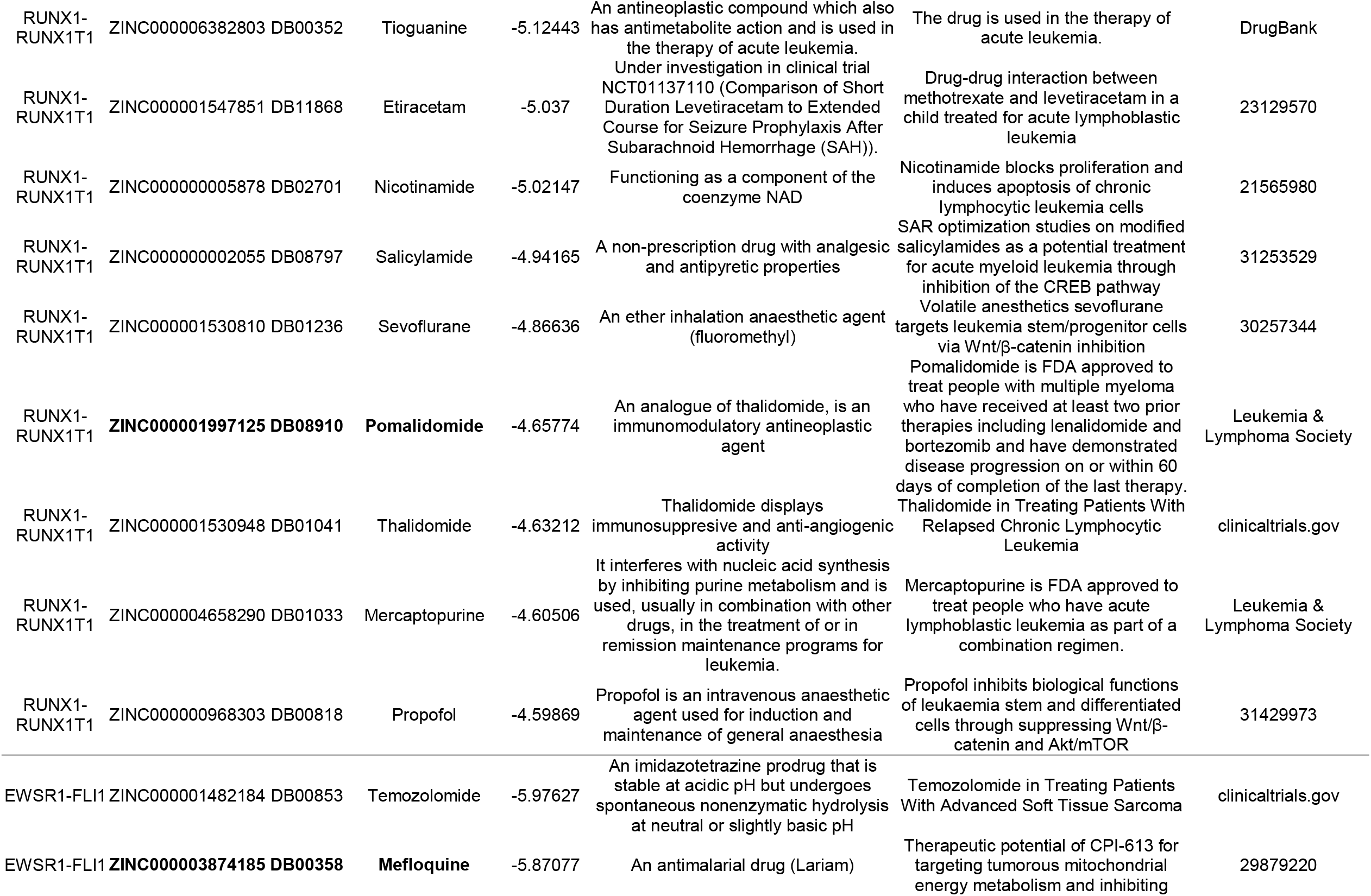
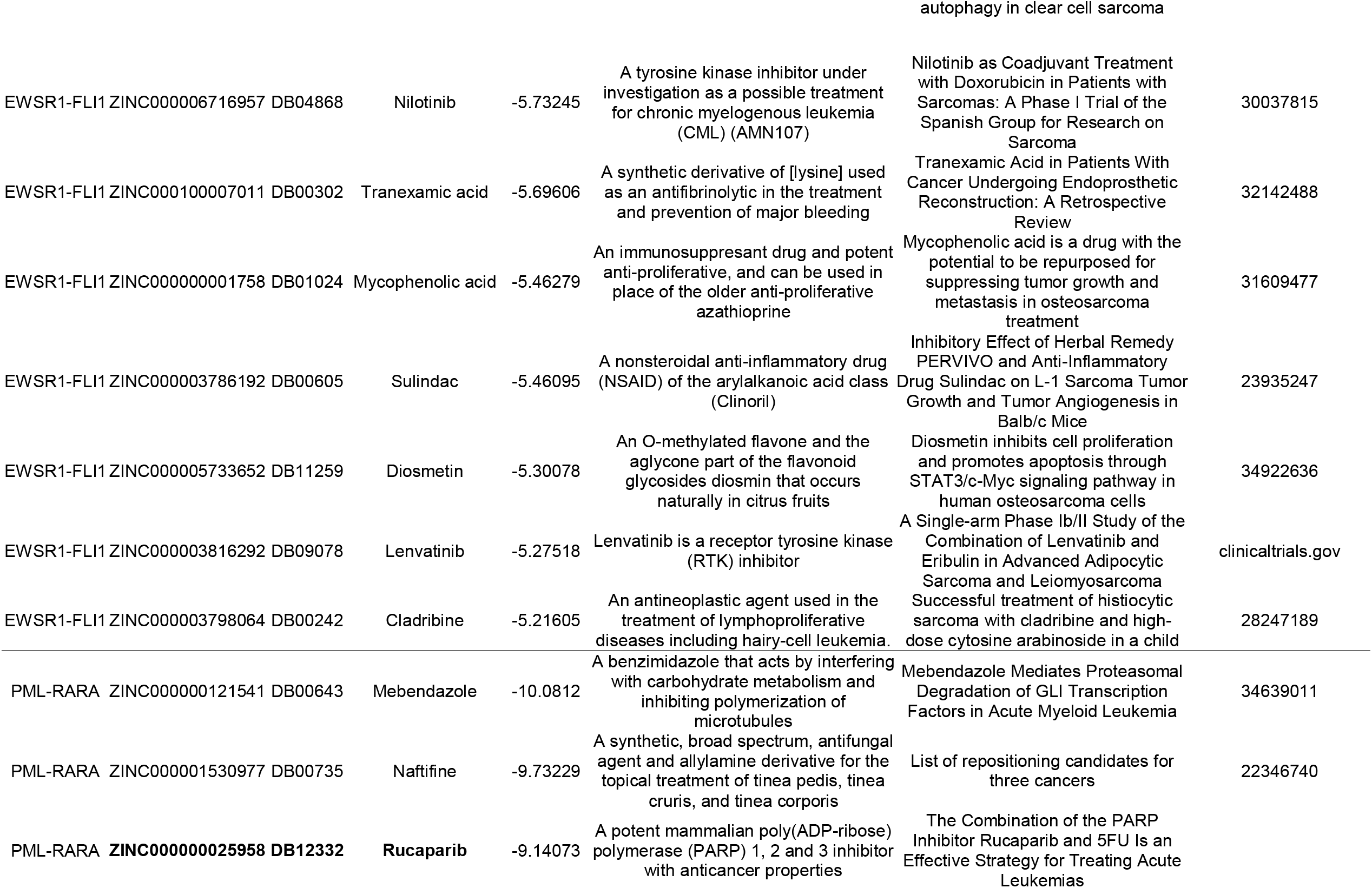
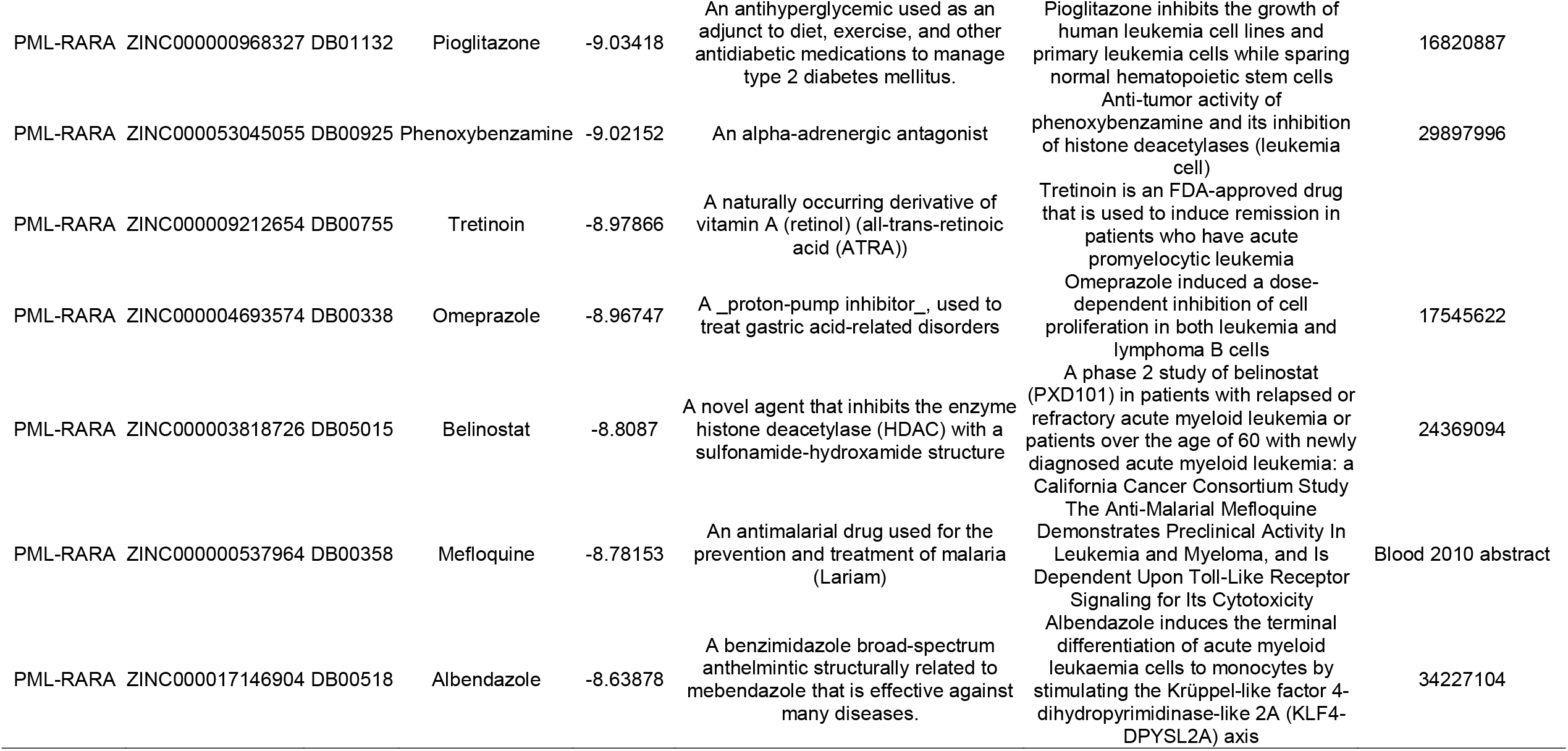
Top 10 individual cancer type-associated FDA-approved drugs of the top five most frequent TFFGs.

### Potential drug candidates of the top five most frequent TFFPs

For the top five most frequent TFFPs, we performed virtual screening in two different strategies. First, we screened the FDA-approved drugs using the predicted active sites by SiteMap (Supplementary Table S3A). Second, we screened the FDA-approved drugs using the known biological domains as the active sites (Supplementary Table S3B).

#### 1. TMPRSS2-ERG

The docking scores of the top 10 interacting FDA-approved drugs interacting with the predicted active sites ranged between -7.41596 and -6.56428 (Supplementary Table S3A). On the contrary, for the screening between TMPTRS2-ERG and known TF domains of ERG versus the FDA-approved drugs, the docking scores of the top 10 drugs ranged between -5.91515 and -5.15658. Only chlordiazepoxide was commonly listed in these two lists with strong binding in the first screening based on the whole 3D structure (-7.0919 < -5.73403). Among the top ten cancer-context-based drugs, we chose the receptor tyrosine kinase inhibitor, sunitinib, which is under clinical trial (Phase II) in men with advanced prostate cancer, as the potential drug repurposing for TMPRSS2-ERG patient^11^. Sunitinib is a receptor tyrosine kinase inhibitor and chemotherapeutic agent used for the treatment of renal cell carcinoma (RCC) and imatinib-resistant gastrointestinal stromal tumor (GIST), according to the explanation of the DrugBank. In the 50 ns run of the simulation, the RMSD plot of the protein-ligand complex showed a stable pattern (Supplementary Figure S1). Sunitinib showed higher interaction in our 3D structure based predicted active sites (-7.15136 > -2.24803).

#### 2. KMT2A-AFF1

The docking score of the top ten drug molecules against the KMT2A-AFF1 ranged between -6.87 and -5.94 from our first screening between our predicted active sites and FDA-approved drugs (Table 1). On the other hand, the second screening between the known TF domains and FDA-approved drugs showed weaker binding (the docking score ranged between -4.77396 and -4.34921). an antineoplastic antibiotic, mitomycin is under clinical testing for patients with acute myelogenous leukemia. We performed an MDS analysis between KMT2A-AFF1 and mitomycin (ZINC000030726187). In the 50 ns run of the simulation, the RMSD plot of the protein-ligand complex showed a stable pattern (Supplementary Figure S2).

#### 3. RUNX1-RUNX1T1

The docking scores in the first screening ranged from -5.91 to -5.11. One of the top 10 drugs. The docking scores in the second screening ranged from -4.77 to -4.34. Chlordiazepoxide and Lysteda were common in both lists. We chose pomalidomide, which is an analog of thalidomide and an immunomodulatory antineoplastic agent and is approved by the FDA to treat people with multiple myeloma. The MDS analysis between RUNX1-RUNX1T1 and pomalidomide (ZINC000001997125) a stable pattern interaction (Supplementary Figure S2).

#### 4. PML-RARA

The combination of the PARP inhibitor rucaparib and 5FU is the currently used strategy for treating acute leukemias. We chose Rucaparib as the candidate for drug repurposing for the PML-RARA fusion-positive patients. In the 50 ns run of the MDS, the RMSD plot of the protein-ligand complex showed a consistent pattern (Supplementary Figure S4). The docking scores in the first screening of the PML-RARA and FDA-approved drugs were ranging from -10.08 to -9.14 (Supplementary Table S3), which is lower than the ones of the top ten molecules in the second screening (from -7.11 to -6.00).

#### 5. EWSR1-FLI1

Out of the top 10 drugs, we chose mefloquine as the potential candidate for drug repurposing for the EWSR1-FLI1 fusion patients. There was a report that mefloquine induced cell death in prostate cancer cells ^12^. The MDS analysis result showed stable interaction between fusion protein and drug (Supplementary Figure S5). The docking scores of the top ten drugs in the first screening ranged from – 6.32 to - 5.87, but the one in the second screening showed weaker interaction (from -4.38 to -3.60). Recently, there was one reported molecule, TK216 under investigation in the clinical trial for EWSR1-FLI protein patients. When we performed the docking experiment with the same active site against the TK216, then it showed a relatively weaker interaction (-4.04) than our screened molecules.

## DISCUSSION

In this study, We tried to develop a computational pipeline of the drug development of transcription factor fusion proteins. Since we didn’t validate the 3D structures (i.e., protein X-ray crystallography), our suggested molecules may not be the best potential repositioning drugs. However, our trial may be worthy as the trigger to advance the development of diver mechanism-related drugs in human cancer fusion genes. Further interpretation following the steps described in this study with examples of the top five most frequent fusion proteins to other TFFPs based on our systematic prediction of whole 3D structures and virtually screened list, we can identify potential repurposing candidates. Our study can be helpful for the systematic view of the human TF fusion proteins and the development of new therapeutic targets and drugs.

## MATERIALS AND METHODS

### Fusion gene information

We obtained fusion gene information from ChimerKB4.0, which is the integration of the manually curated fusion genes from Mitelman DB, COSMIC, and GenBank as a subcategory of ChimerDB 4.0 fusion gene knowledgebase ^4^. We obtained 1597 fusion genes with publication support and experimental evidence. Out of these 1007 fusion genes had the breakpoint information. After checking the open reading frames, only 266 fusion genes were in-frame and potentially make the fusion proteins. Out of these 266 in-frame fusion genes, 127 were transcription factor fusion genes. In this study, we focused on these 127 TFFGs.

### Open reading frame (ORF) annotations

Between the 5’-partner gene and the 3’-partner gene, we checked the open reading frame of the full-length fusion transcript sequence. When both breakpoints of 5’- and 3’-genes are located inside of coding region (CDS) and the number of fusion transcript sequences from the transcription start site of 5’-gene to the transcription end site of 3’-gene is a multiple of three, then we reported this fusion gene as ‘in-frame’. If there are one or two nucleotide insertions, then we reported it as the ‘frame-shift’. Except for these two ORFs, there are 15 more ORFs such as ‘3UTR-CDS’, ‘3UTR-3UTR’, ‘3UTR-5UTR’, ‘3UTR-intron’, ‘CDS-3UTR’, ‘CDS-5UTR’, ‘CDS-intron’, ‘5UTR-CDS’, ‘5UTR-3UTR’, ‘5UTR-5UTR’, ‘5UTR-intron’, ‘intron-CDS’, ‘intron-3UTR’, ‘intron-5UTR’, and ‘intron-intron’. Here, the ‘intron’ is reported when the breakpoint is located 6bp apart from the exon junction site to the intron direction. Since our fusion breakpoints were derived from the ESTs and RNA-seq data, all the breakpoints should be located inside the exon. Therefore, if the breakpoint is located on the intron, then we report it as an intron, and the ORFs including the intron in at least one of the partners set aside these categories to not available (NA) ORF cases in our ORF classification. For these analyses, we considered all matched Ensembl transcripts (ENSTs) ^13^. For the 127 in-frame TFFGs, we created the full-length fusion transcript sequences and we ran the open reading frame finder (ORFfinder), which is mainly selecting the longest ORFs from the six-frames-based translation ^14^. Then, we had 732 TF fusion amino acid sequences.

### Retention analysis of 39 protein features from UniProt

We first downloaded the GFF (General Feature Format) format protein information of UniProt accessions from UniProt for the genes involved in 127 fusion genes ^15^. UniProt provides the loci information of 39 protein features including six molecule processing features, 13 region features, four site features, six amino acid modification features, two natural variation features, five experimental info features, and three secondary structure features. Since such feature loci information was based on amino acid sequence, the genomic breakpoint information was converted into the amino acid level while considering all UniProt protein accessions, ENST isoforms, and multiple breakpoints for each partner. To map each feature to the human genome sequence, we used the GENCODE gene model of the human reference genome v19 ^16^. For the 5’-partner gene, we considered the protein feature to be retained in the fusion gene if the breakpoints occurred on the 3’-end of the protein feature. On the contrary, if a protein domain was not included completely in the fusion amino acid sequence, we reported that such fusion genes did not retain that protein feature. Similarly, for the 3’-partner gene, we considered the fusion gene to have retained the protein feature if the breakpoints occurred on the 5’-end of the protein feature region. For TFFGs, we checked the retention and ORFs of the main protein functional features that may include the DNA binding features, such as ‘DNA binding’, ‘domain’, ‘motif’, ‘region’, and ‘zinc finger’.

### Fusion protein sequence information and 3D structure prediction of fusion proteins

The 3D structures were adopted from our recent work, FusionPDB, which is available from https://compbio.uth.edu/FusionPDB. The approaches are as follows. We used the GPU-based Linux version AlphaFold2 following the step-by-step procedures in the GitHub repository of AlphaFold2. We input the fasta format files for all 732 TF fusion protein’s amino acids into AlphaFold2. For the alignment, we have downloaded the databases (size: 2.2 TB) such as uniref90, uniport, uniclust30, small_bfd, pdb_seqres, pdb_mmcif, pdb70, params, mgnify, and bfd. The model parameters were made available by Alphafold2 for users. The output folder provided the predicted PDB structures on the basis of model confidence and ranked as ranked_0.pdb to ranked_4.pdb.

### Preparation of protein and active site identification

The PDB structures of predicted proteins may contain missing atoms, missing connectivity information, and water molecules, which need to be removed or preprocessed before using the 3D structure of the protein for virtual screening. All 1267 predicted protein structures were preprocessed using the protein preparation module of Schrödinger (Release 2022-2, LLC, New York, NY, 2021) ^17^, by removing the water molecules, assigning the bond orders, and adding the hydrogen bonds to the crystal structure of receptor molecule. Restrained minimization was done by the constraint of the RMSD and OPLS force field. To identify the active sites of all 732 predicted TF fusion proteins, we used the SiteMap of Schrödinger (release 2021-4). The SiteMap provides the site score for the predicted active site on the basis of multiple factors such as the size of the site, exposure to the solvent, hydrophobicity, hydrophilicity, and hydrogen bond donor and acceptor.

### Known DNA binding and active domains

For the top five most frequently involved TFs, we downloaded the DNA binding and active domain residue information from the protein functional feature annotation of UniProt.

### Receptor grid generation and Virtual screening

The grid size represents the volume of a receptor’s active sites where the ligand can search for binding while docking. The grid around the receptor was generated using a module GLIDE available with the Glide of Schrödinger package ^18^. The dimensions of the grid were selected by considering the active sites predicted by SiteMap. Virtual screening is a computational technique used in the drug discovery process to get small molecules that are most likely to bind to receptors or target molecules ^19^. In this work, we considered all FDA-approved ligand libraries of the ZINC database accessed on 10^th^ December 2021. These selected libraries were processed by the LigPrep of Schrödinger package and made available for virtual screening. In silico screening and docking analysis of 732 predicted TF fusion protein targets against the FDA-approved ligands were carried out using the Glide of Schrödinger package (release 2021-4) ^20^. We also tried to investigate some new ligands that can interact with the same active site. For this, we made a ligand library with 320 K compounds which consists of natural, endogenous, metabolites, in-trials, and in-vivo-only molecules.

### Molecular dynamics simulation (MDS)

MDS was used to analyze the physical movement of atoms and molecules during drug target and ligand interaction in the system. In this study, we selected the top five most frequent fusion proteins and used Desmond of the Schrödinger package (release 2021-4) for MDS analysis. The coordinates for MD simulation were obtained from the receptor-ligand complex. The system builder panel was used to prepare the complex for simulation by selecting the solvation model TIP3P. Boundary conditions were set as an orthorhombic box with the dimensions of 10 Å × 10 Å × 10 Å, to ensure that the entirety of the complex was covered in the solvent. The system was neutralized by balancing the net charge of the system by adding three Cl-counter ions. The system was minimized for 2000 iterations of a hybrid of the steepest descent and the limited memory Broyden-Fletcher-Gold farb-Shanno (LBFGS) algorithms, with a convergence threshold of 1.0 kcal/mol/Å. The whole system was subjected to 300 K and 1.0325 bar pressure, for 50 ns at NPT ensemble of MD simulation with a recording interval of 50 ps for total energy (kcal/mol). The structural modifications and real-time behavior of the complex structure were computed by means of the root mean square deviation (RMSD) and potential energy of the receptor-ligand complex during the course of 50 ns simulation time ^21^.

## ACKNOWLEDGEMENTS

We thank the members of the Center for Computational Systems Medicine at SBMI at UTHealth and Dr. Joseph A Ludwig at the MD Anderson Cancer Center for valuable questions, discussions, and suggestions.

## DATA AVAILABILITY

Transcription factor fusion proteins’ sequence, structure, and interacting small molecule information are available from the FusionPDB website (https://compbio.uth.edu/FusionPDB). Further information and requests should be directed to Dr. Pora Kim.

## FUNDING

This work was partially supported by the National Institutes of Health grants [R35GM138184] to P. Kim. The funders had no role in study design, data collection, and analysis, decision to publish, or preparation of the manuscript. Funding for open access charge: Startup Fund to Dr. Kim from the University of Texas Health Science Center at Houston.

### Conflict of interest statement

None declared.

## Key points

- Our study provides the whole 3D structures of 732 transcription factor fusion proteins.
- Our study provides a pipeline to predict the potentially interacting small molecules with TF fusion proteins.
- Our study provides potential drug repurposing of TMPRSS2-ERG, KMT2A-AFF1, RUNX1-RUNX1T1, PML-RARA, and EWSR1-FLI1.

## TABLES AND FIGURES LEGENDS

**Figure 5.**
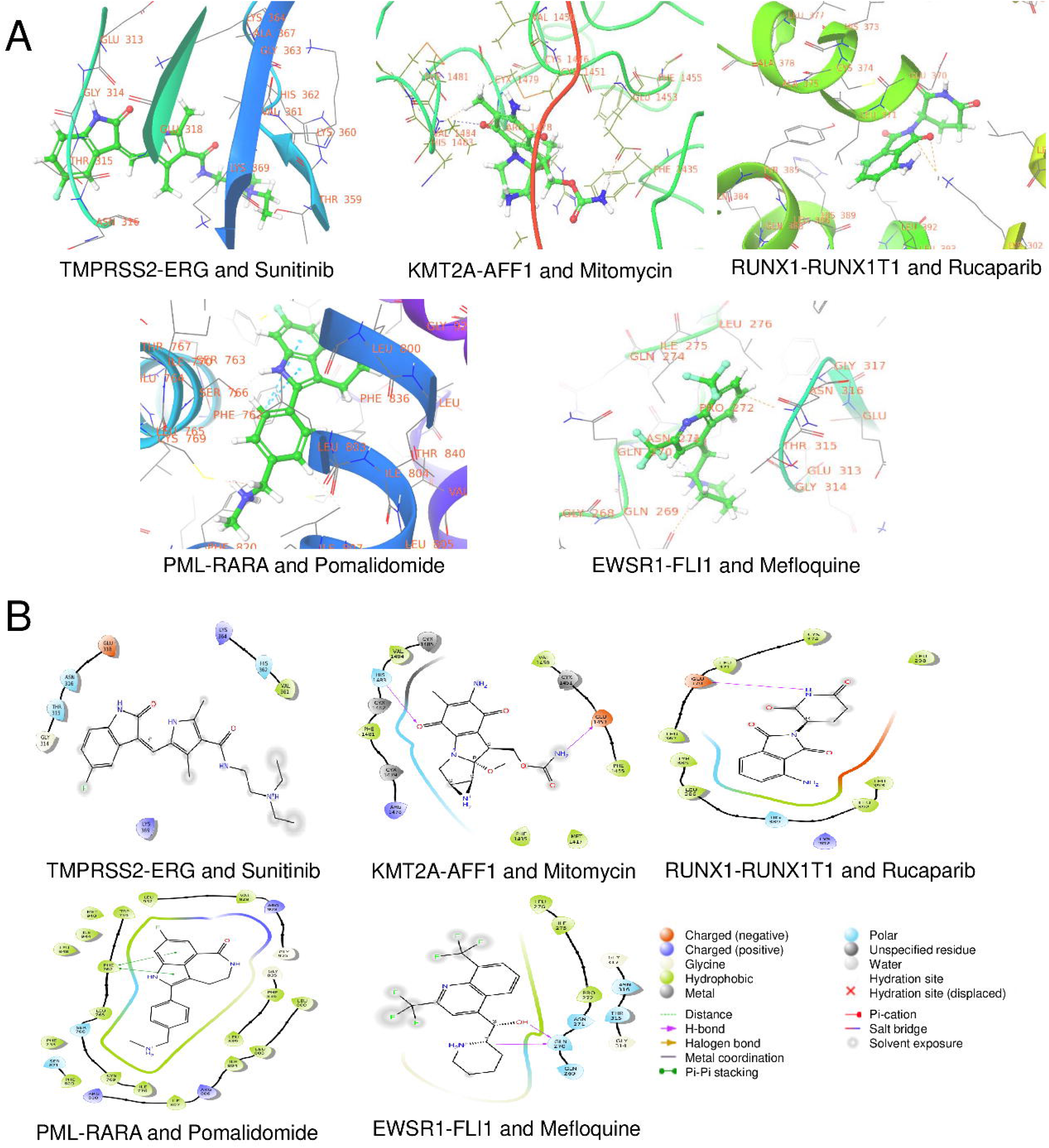
Top five most frequent TFFPs and chosen FDA-approved small molecule complex in 3D. (A) TMPRSS2-ERG and sunitinib. (B) KMT2A-AFF1 and Mitomycin. (C) RUNX1-RUNX1T1 and Pomalidomide. (D) PML-RARA and rucaparib. (E) EWSR1-FLI1 and mefloquine.

**Supplementary Table S1**. TFFP information.

**Supplementary Table S2**. Statistics of the TF and target genes in TF fusion genes.

**Supplementary Table S3**. Top 10 FDA-approved drugs interacting with TF fusion proteins based on different active sites. (A) Top 10 FDA-approved drugs interacting with 5 TF fusion proteins based on the predicted active sites by the SiteMap based on the whole 3D fusion protein structure. (B) Top 10 FDA-approved drugs interacting with 5 TF fusion proteins based on the biological active sites (DNA binding domain and active domain). (C) Comparison of the docking scores of the chosen drugs that have good MDS results based on the predicted active sites by the SiteMap and based on the biological active sites (DBD and AD).

**Supplementary Figure S1**. MDS analysis result between TMPRSS2-ERG and sunitinib.

**Supplementary Figure S2**. MDS analysis result between KMT2A-AFF1 and Mitomycin.

**Supplementary Figure S3**. MDS analysis result between RUNX1-RUNX1T1 and Pomalidomide.

**Supplementary Figure S4**. MDS analysis result between PML-RARA and Rucaparib.

**Supplementary Figure S5**. MDS analysis result between EWSR1-FLI1 and Mefloquine.

## Notes

### Competing Interest Statement

The authors have declared no competing interest.

## REFERENCES

1 Kim, P., Yiya, K. & Zhou, X. FGviewer: an online visualization tool for functional features of human fusion genes. Nucleic Acids Res 48, W313–W320, doi:10.1093/nar/gkaa364 (2020).

2 Graff, R. E. et al. The TMPRSS2:ERG fusion and response to androgen deprivation therapy for prostate cancer. Prostate 75, 897–906, doi:10.1002/pros.22973 (2015).

3 Zollner, S. K. et al. Inhibition of the oncogenic fusion protein EWS-FLI1 causes G2-M cell cycle arrest and enhanced vincristine sensitivity in Ewing’s sarcoma. Sci Signal 10, pdoi:10.1126/scisignal.aam8429 (2017).

4 Jang, Y. E. et al. ChimerDB 4.0: an updated and expanded database of fusion genes. Nucleic Acids Res 48, D817–D824, doi:10.1093/nar/gkz1013 (2020).

5 Jumper, J. et al. Highly accurate protein structure prediction with AlphaFold. Nature 596, 583–589, pdoi:10.1038/s41586-021-03819-2 (2021).

6 UniProt, C. UniProt: the universal protein knowledgebase in 2021. Nucleic Acids Res 49, D480–D489, doi:10.1093/nar/gkaa1100 (2021).

7 Han, H. et al. TRRUST v2: an expanded reference database of human and mouse transcriptional regulatory interactions. Nucleic Acids Res 46, D380–D386, doi:10.1093/nar/gkx1013 (2018).

8 Rose, P. W. et al. The RCSB Protein Data Bank: views of structural biology for basic and applied research and education. Nucleic Acids Res 43, D345–356, doi:10.1093/nar/gku1214 (2015).

9 Wishart, D. S. et al. DrugBank 5.0: a major update to the DrugBank database for 2018. Nucleic Acids Res 46, D1074–D1082, doi:10.1093/nar/gkx1037 (2018).

10 Sastry, G. M., Adzhigirey, M., Day, T., Annabhimoju, R. & Sherman, W. Protein and ligand preparation: parameters, protocols, and influence on virtual screening enrichments. J Comput Aided Mol Des 27, 221–234, doi:10.1007/s10822-013-9644-8 (2013).

11 Dror Michaelson, M. et al. Phase II study of sunitinib in men with advanced prostate cancer. Ann Oncol 20, 913–920, doi:10.1093/annonc/mdp111 (2009).

12 Yan, K. H. et al. Mefloquine induces cell death in prostate cancer cells and provides a potential novel treatment strategy in vivo. Oncol Lett 5, 1567–1571, doi:10.3892/ol.2013.1259 (2013).

13 Howe, K. L. et al. Ensembl 2021. Nucleic acids research 49, D884–D891, doi:10.1093/nar/gkaa942 (2020).

14 Sayers, E. W. et al. Database resources of the National Center for Biotechnology Information in 2023. Nucleic Acids Res, doi:10.1093/nar/gkac1032 (2022).

15 UniProt Consortium, T. UniProt: the universal protein knowledgebase. Nucleic acids research 46, 2699, doi:10.1093/nar/gky092 (2018).

16 Harrow, J. et al. GENCODE: the reference human genome annotation for The ENCODE Project. Genome research 22, 1760–1774, doi:10.1101/gr.135350.111 (2012).

17 Madhavi Sastry, G., Adzhigirey, M., Day, T., Annabhimoju, R. & Sherman, W. Protein and ligand preparation: parameters, protocols, and influence on virtual screening enrichments. Journal of computer-aided molecular design 27, 221–234 (2013).

18 Halgren, T. A. et al. Glide: a new approach for rapid, accurate docking and scoring. 2. Enrichment factors in database screening. Journal of medicinal chemistry 47, 1750–1759 (2004).

19 Rester, U. From virtuality to reality-Virtual screening in lead discovery and lead optimization: a medicinal chemistry perspective. Current opinion in drug discovery & development 11, 559–568 (2008).

20 Friesner, R. A. et al. Glide: a new approach for rapid, accurate docking and scoring. 1. Method and assessment of docking accuracy. Journal of medicinal chemistry 47, 1739–1749 (2004).

21 Jacobson, M. P., Friesner, R. A., Xiang, Z. & Honig, B. On the role of the crystal environment in determining protein side-chain conformations. Journal of molecular biology 320, 597–608 (2002).

