## Supplementary figures and images for "Computational design of DNA binding domain-retained fusion proteins and virtual screening against FDA-approved drugs"

### Supplemental Figure 1

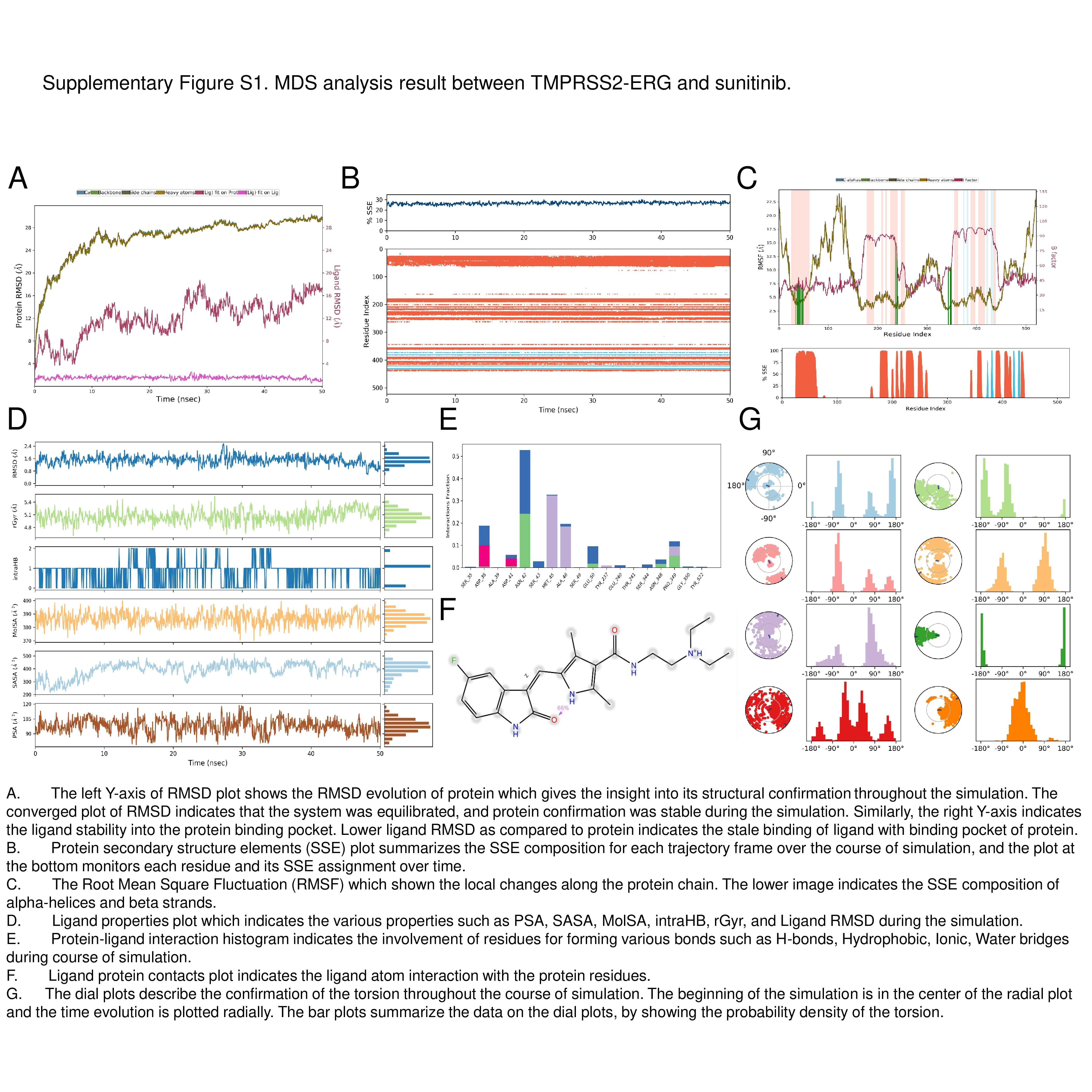

### Supplemental Figure 2

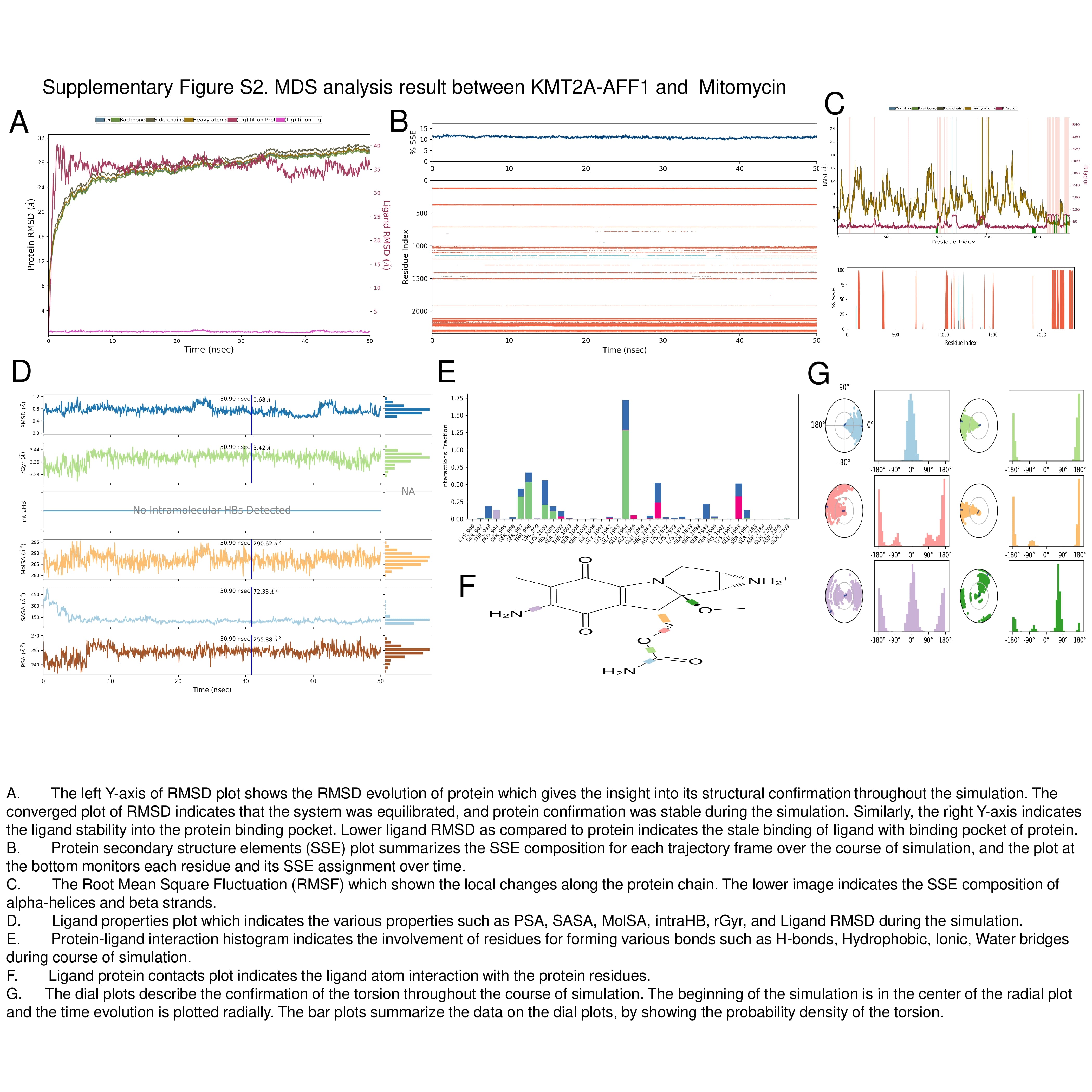

### Supplemental Figure 3

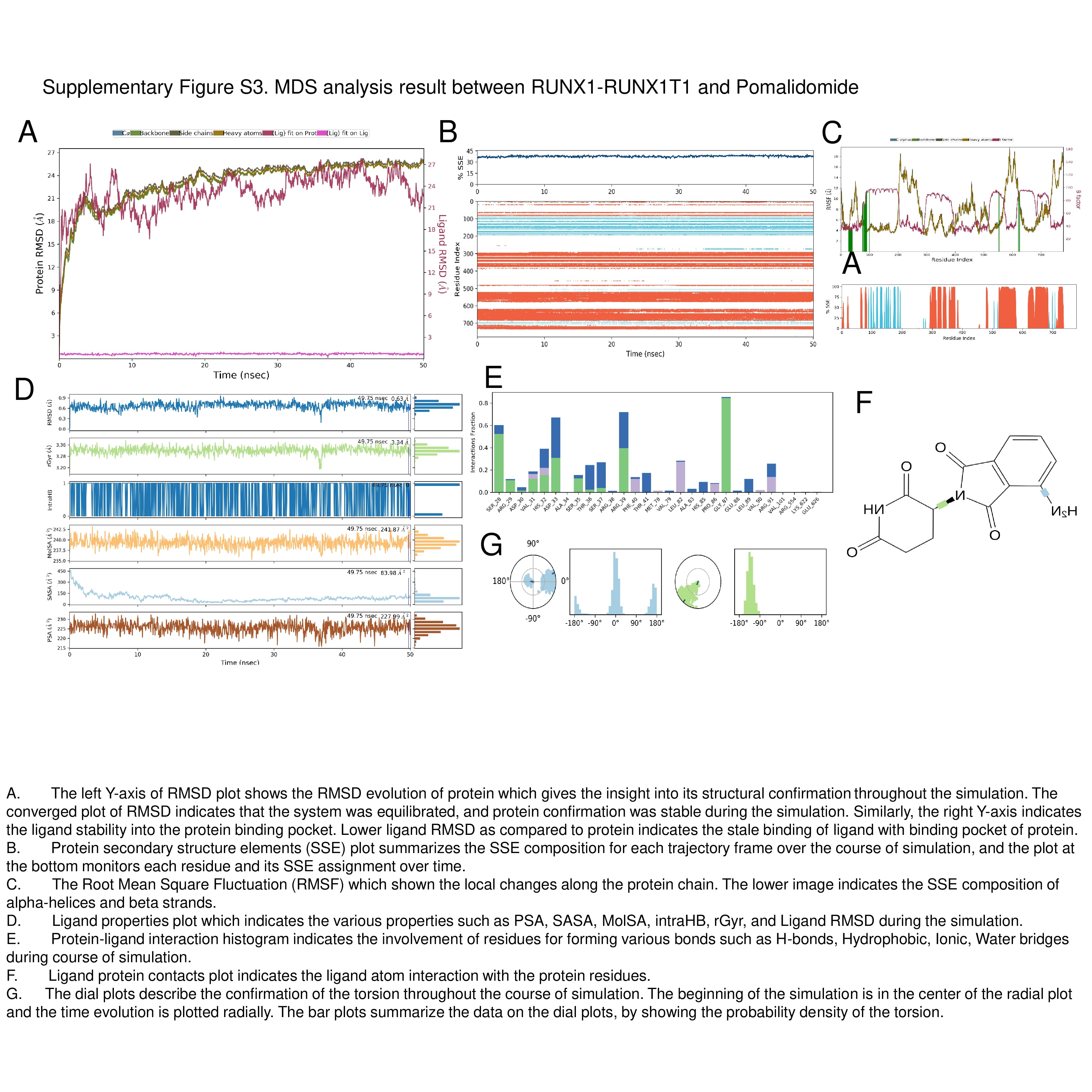

### Supplemental Figure 4

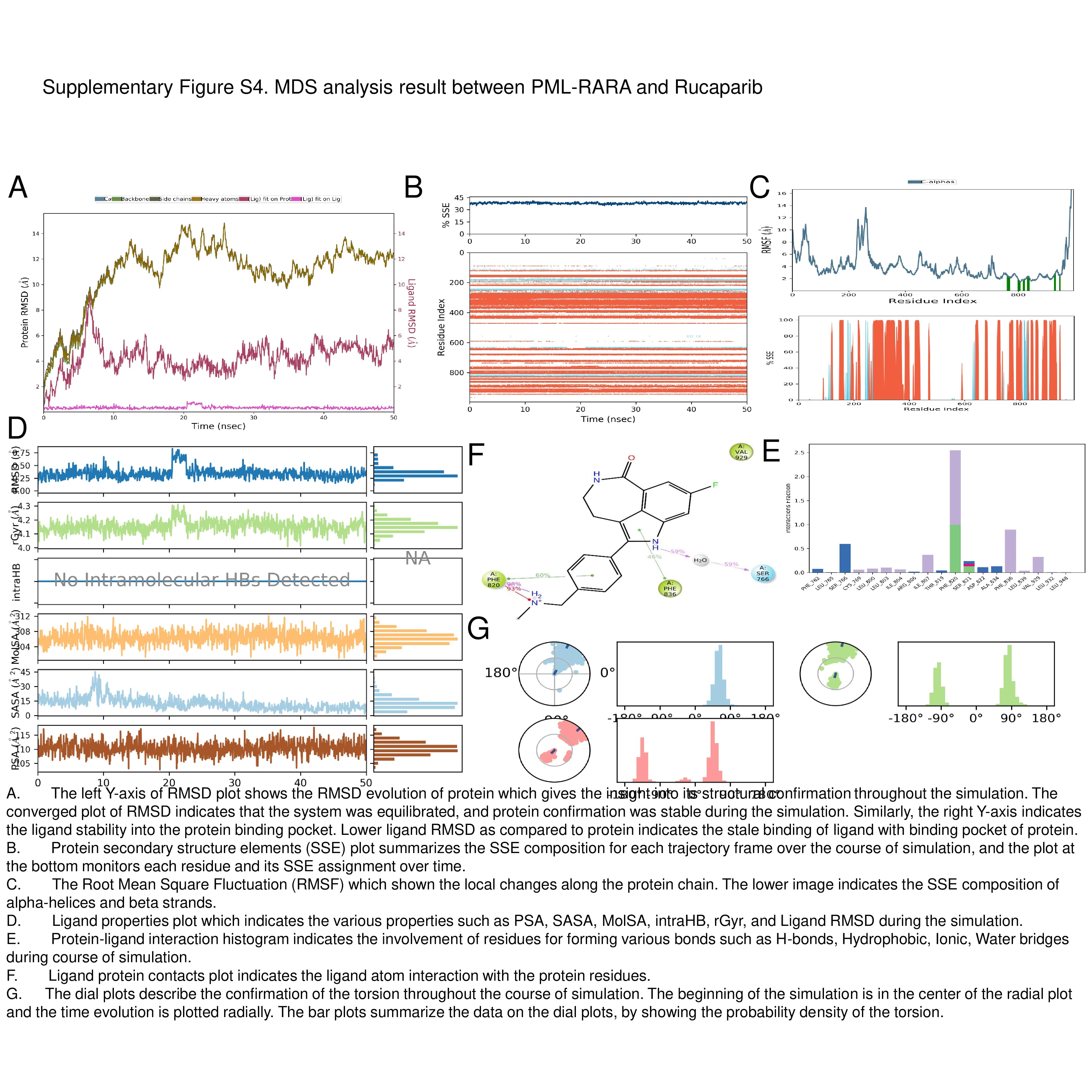

### Supplemental Figure 5

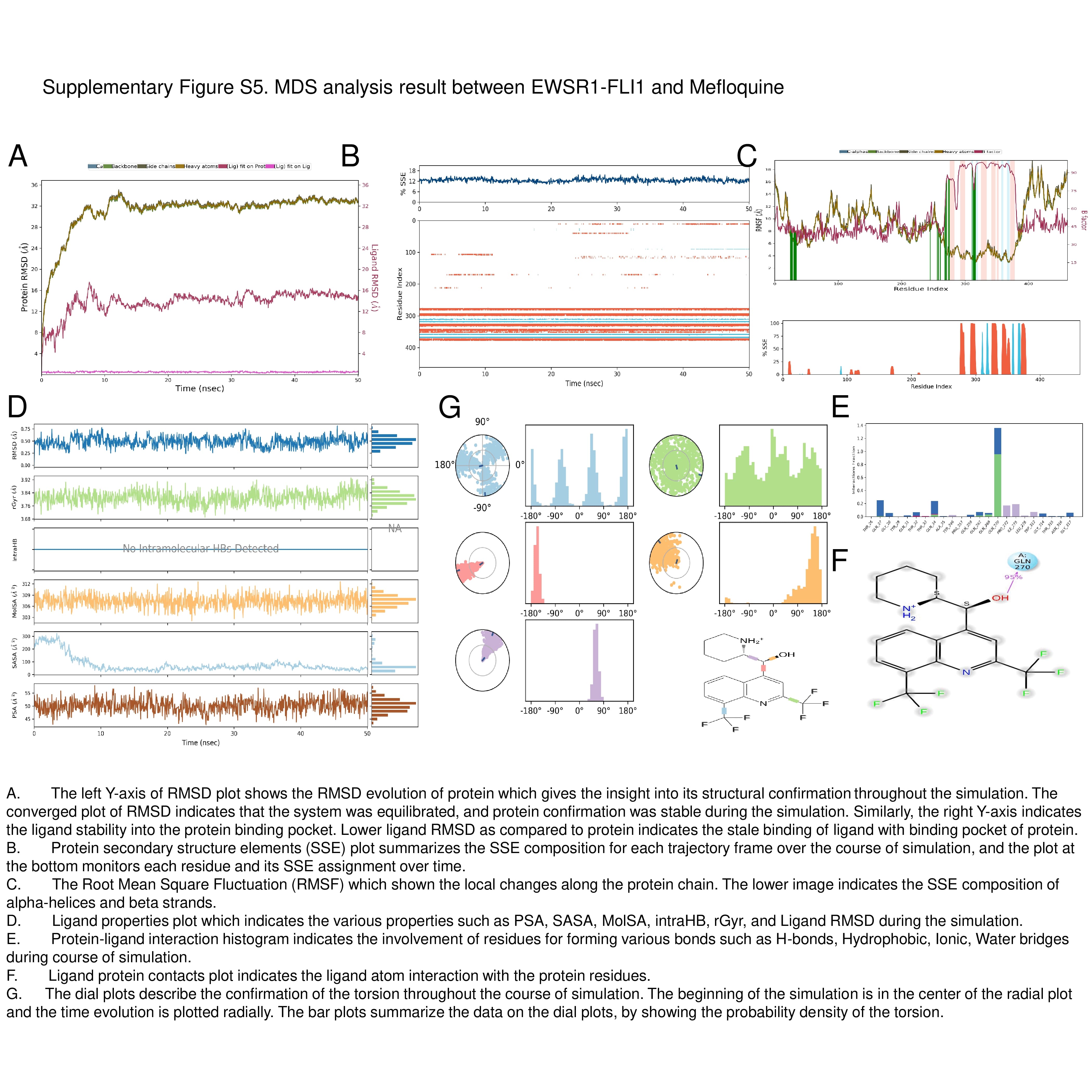
